# A functional microbiome catalog crowdsourced from North American rivers

**DOI:** 10.1101/2023.07.22.550117

**Authors:** Mikayla A. Borton, Bridget B. McGivern, Kathryn R. Willi, Ben J. Woodcroft, Annika C. Mosier, Derick M. Singleton, Ted Bambakidis, Aaron Pelly, Filipe Liu, Janaka N. Edirisinghe, José P. Faria, Ikaia Leleiwi, Rebecca A. Daly, Amy E. Goldman, Michael J. Wilkins, Ed K. Hall, Christa Pennacchio, Simon Roux, Emiley A. Eloe-Fadrosh, Stephen P. Good, Matthew B. Sullivan, Christopher S. Henry, Elisha M. Wood-Charlson, Matthew R.V. Ross, Christopher S. Miller, Byron C. Crump, James C. Stegen, Kelly C. Wrighton

## Abstract

Predicting elemental cycles and maintaining water quality under increasing anthropogenic influence requires understanding the spatial drivers of river microbiomes. However, the unifying microbial processes governing river biogeochemistry are hindered by a lack of genome-resolved functional insights and sampling across multiple rivers. Here we employed a community science effort to accelerate the sampling, sequencing, and genome-resolved analyses of river microbiomes to create the Genome Resolved Open Watersheds database (GROWdb). This resource profiled the identity, distribution, function, and expression of thousands of microbial genomes across rivers covering 90% of United States watersheds. Specifically, GROWdb encompasses 1,469 microbial species from 27 phyla, including novel lineages from 10 families and 128 genera, and defines the core river microbiome for the first time at genome level. GROWdb analyses coupled to extensive geospatial information revealed local and regional drivers of microbial community structuring, while also presenting a myriad of foundational hypotheses about ecosystem function. Building upon the previously conceived River Continuum Concept^1^, we layer on microbial functional trait expression, which suggests the structure and function of river microbiomes is predictable. We make GROWdb available through various collaborative cyberinfrastructures^2, 3^ so that it can be widely accessed across disciplines for watershed predictive modeling and microbiome-based management practices.

## Main

Earth’s surface is dominated by water, much of it the oceans, which is known to buffer against anthropogenic climate change via microbes dictating the fate of ocean-absorbed carbon^4^. While the oceans and their microbes have been extensively studied globally by large scientific consortia (e.g. *Tara* Oceans Consortium^5^), other elements of Earth’s water system, such as rivers, are relatively understudied. This is problematic as rivers (i) offer an important nexus of nutrient transport across terrestrial and aquatic interfaces^6^, (ii) are hotspots for biogeochemical processes that significantly contribute to global terrestrial carbon and nitrogen budgets ultimately impacting global greenhouse gas (GHG) emissions, eutrophication, and acidification^6–8^, and (iii) have immediate societal impacts on sustainable energy, agriculture, environmental health, and human health^9–11^. Microbial metabolisms dictate river ecosystem functioning with major influence on carbon (C) respiration and sequestration, nitrogen (N) cycling and uptake, food webs, and pollutants (e.g., Hg, As)^12–16^. Given these important contributions there is a growing need to better resolve the ecology, functional potential, and biogeochemical contributions of microbes across diverse river systems.

Despite being critical modulators of biogeochemistry, river microbiomes remain under sampled due to methodological constraints and sampling scope. For example, a majority of river microbiome studies rely only on 16S rRNA gene analysis (**Extended Data File 1**). While these single gene studies have advanced understanding of riverine microbial community diversity and membership^17–20^, they lack information on poorly characterized lineages and are limited in their capacity to functionally link microorganisms to biogeochemical processes. Albeit less, there are several studies with metagenomics that provide functional attributes of river microbiomes, but these rely heavily on existing databases rather than *de novo* assembly and metagenome assembled genomes (**Extended Data File 1**), masking the contributions of novel members of the microbiome. Less than five studies employ genome-resolved expression methods hindering estimates of the metabolic processes that are active in river systems (**Extended Data File 1**). Finally, in terms of sampling, most studies focus on a single site or stream network (**Extended Data File 1**), leaving the generalizability of microbiome rules across river systems uncertain. To establish a transferable functional understanding of river microbiomes, there is a need to genomically resolve the taxonomy, metabolic potential, and expression of river microbiomes at scale.

To meet this need, we developed a crowd-sourced, distributed sampling effort to increase and standardize river microbiome sampling globally, compiling these sequencing results into a large-scale Genome Resolved Open Watersheds database (GROWdb). An emphasis of GROWdb is a publicly available and ever-expanding microbial genome database that has accessible content. GROWdb represents the first microbial, river-focused resource parsed at various scales from genes to MAGs to community level including expression and potential based measurements that will be of interest to microbiologists, ecologists, geochemists, hydrologists, and modelers. GROWdb is based on a crowd-sourced, network-of-networks approach^21^ to move beyond a small collection of well-studied rivers, towards a spatially distributed, global network of systematic observations.

### Construction of the Genome Resolved Open Watersheds database (GROWdb) enabled by community sampling and sequencing

To establish the GROWdb, over 100 teams were crowd-sourced to broadly sample 163 sites across United States rivers and develop ∼3.8 terabases of metagenomic and metatranscriptomic sequencing data to go with extensive (up to 287) geochemical and geospatial measurements at each site (**Fig. 1A**, detailed per-sampling data available in **Fig. 1B** and **Extended Data File 1**). Through this, we aimed to capture community-level, genome-resolved microbiome variations in taxonomy, function, and gene expression in the context of geographic and environmental gradients across the United States. The effort resulted in surface water sampling that covered 90% of United States watersheds (n=21 as determined by hydrologic unit 2) (**Fig. 1C**) and spanned diverse ecoregions, stream orders, and watershed sizes (**Extended Data Fig. 1**). Collectively, GROWdb integrates genomics, biogeochemistry, and a broad range of contextual environmental variables into a platform that enables a predictive framework of microbiomes and their biogeochemical contributions.

**Figure 1.**
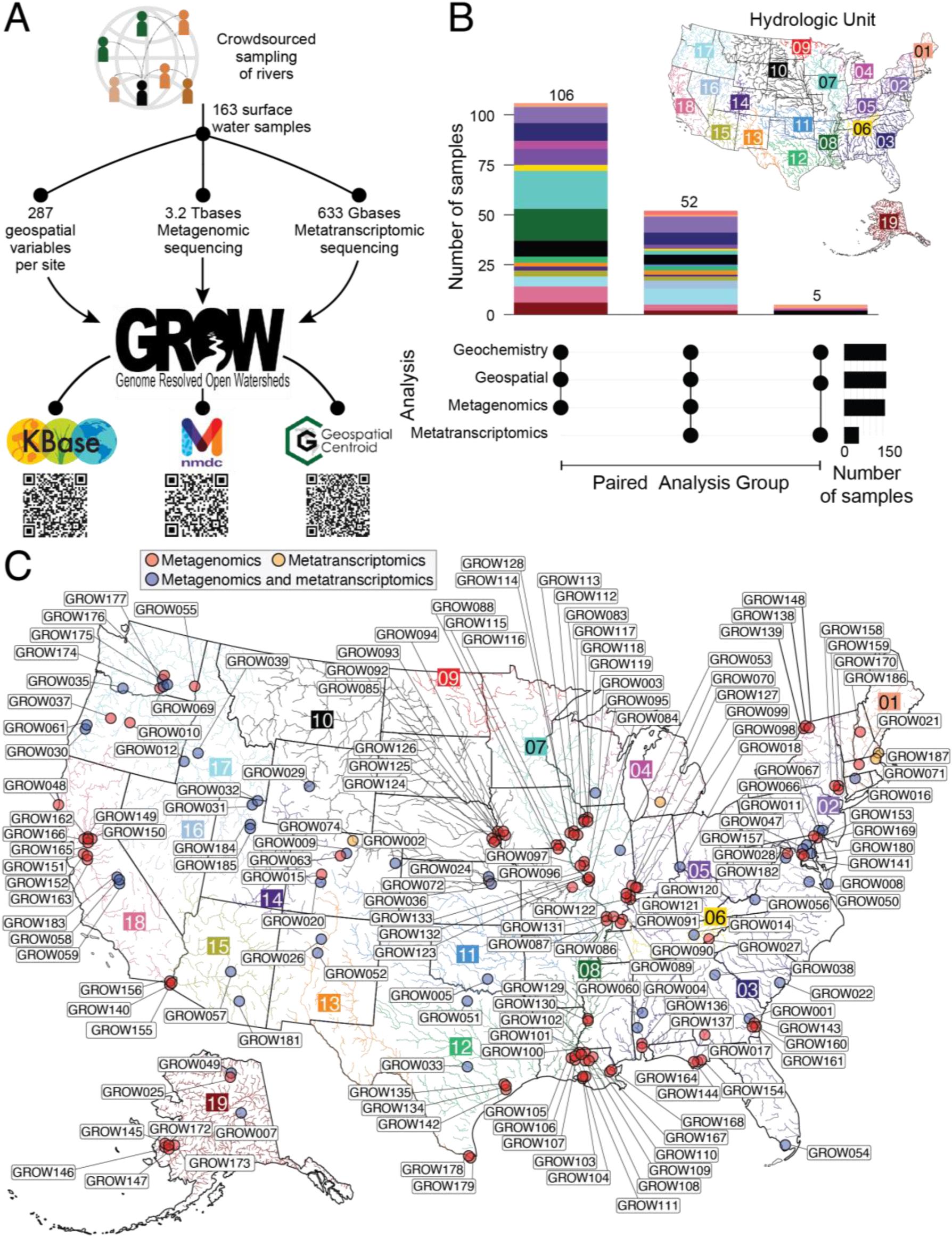
Distributed sampling and sequencing of rivers enabled the construction of the Genome Resolved Open Watersheds (GROW). A) Workflow denoting the number of samples and resulting datasets made up of geospatial and microbiome (metagenomics, metatranscriptomics) data. GROWdb data are accessible through KBase, NMDC, and the Geospatial Centroid, with QR codes directing to online data location. B) Plot showing the number of samples with paired data types (denoted as filled black circles below) as stacked bars colored by hydrologic unit, and number of samples per analysis as black solid bars. C) GROW sampling across the United States with points marking a sampling location, color-coded by the microbiome analysis performed (MetaG, red; MetaT, yellow; paired MetaG and MetaT, blue). Boxed numbers and corresponding river color indicate hydrologic unit (HUC-2).

To ensure data accessibility we provide four access points for user engagement with GROWdb (**Fig. 1A**). First, all reads and MAGs are publicly hosted on National Center for Biotechnology (NCBI), enabling transferability to resources that pull and incorporate this content (e.g., Genome Taxonomy Database^22^). Datasets underlying GROWdb are freely available and searchable through the National Microbiome Data Collaborative (NMDC)^23^ data portal, linking to other data types (e.g., metabolome^24^) to allow for broader synthesis where available. GROWdb MAGs are available as an annotated genomic collection in the freely accessible KBase^3^ cyberinfrastructure. Here users can access sample information, gene- and MAG-level annotations, profile functional summaries, and genome scale models in a point and click interface. Lastly, to aid in data exploration we distilled the taxonomic and functional insights from GROWdb in a web accessible format, called GROWdb Explorer, allowing the rapid profiling of taxonomic and functional distributions across the data set. GROWdb version 1 can be accessed across platforms (**Fig. 1A**, QR codes), making this microbiome content available in an expanding repository to incorporate and unify global river multi-omic data for the future.

### Over 2,000 unique microbial genomes recovered from United States surface waters

To uncover the key microbial players and functions in surface water river microbiomes, we constructed a genome database composed of metagenome-assembled genomes (MAGs). Our sequencing represents on average of 3-fold more sequencing per sample compared to published riverine metagenome studies^25^, thereby increasing the sensitivity for detecting the breadth of microbial functions encoded in these systems (**Extended Data Fig. 2**). From this sequencing, we assembled and reconstructed 3,825 medium and high-quality MAGs, which were dereplicated into 2,093 unique MAGs at 99% identity (**Fig. 2, Extended Data File 2**). Based on read mapping, the majority (mean=52%) of metagenomic reads mapped back to this surface water derived MAG database, signifying that the underlying sequencing reads were well represented by the genomic database.

**Figure 2:**
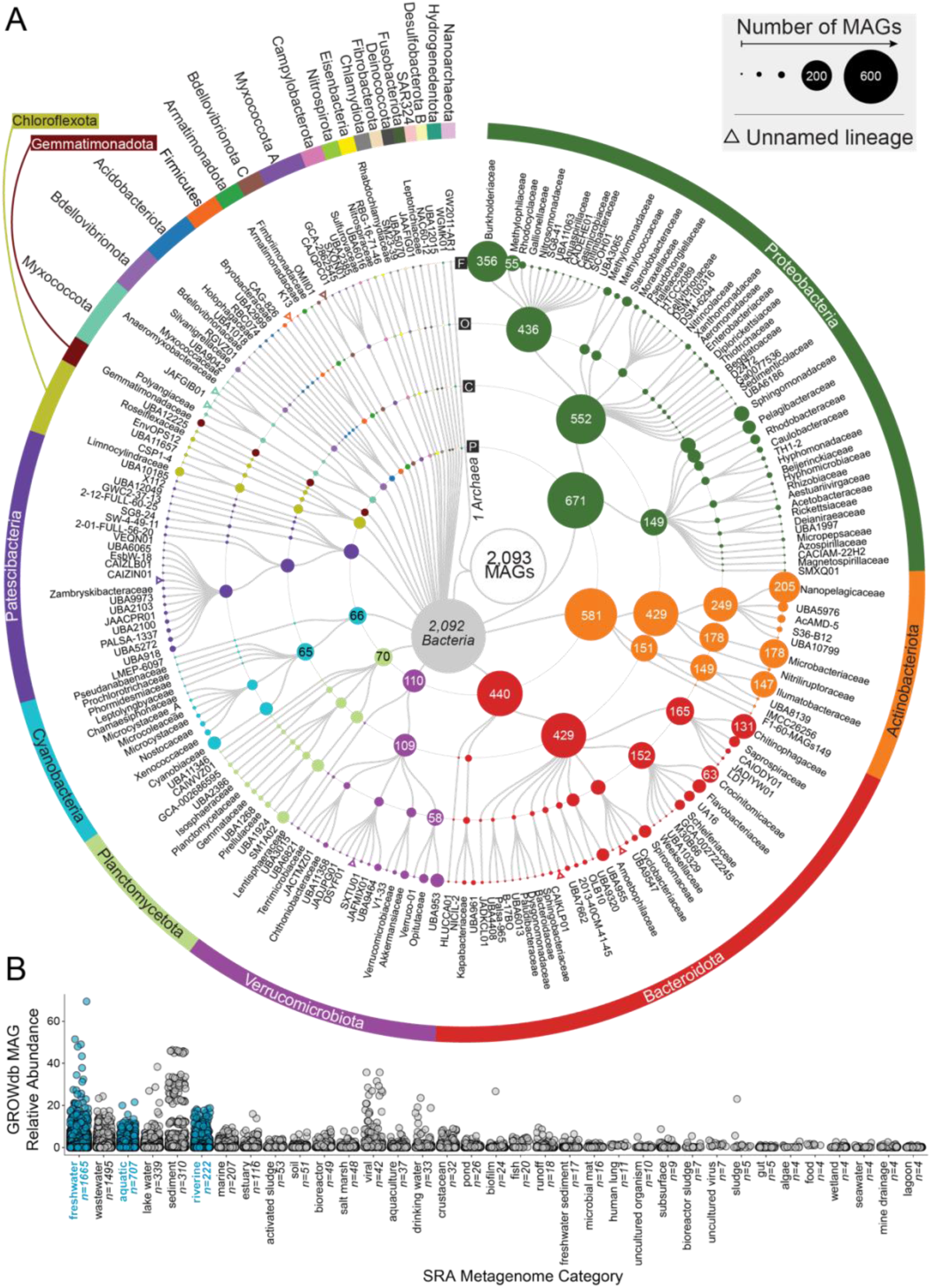
Taxonomic diversity of 2,093 unique surface water metagenome assembled genomes (MAGs) in GROWdb. A) Cladogram shows GROWdb MAGs taxonomy with each sequential ring noting taxonomy level (Phylum, P; Class, C; Order, O; Family, F). Circle size indicates the number of genomes within a given taxonomy level and is further noted by MAG number inside the circle when sampling at that taxonomic position exceeds 50 MAGs sampled. Colors highlight phylum level taxonomy denoted on the outermost ring. Open triangles represent unnamed lineages within a particular level of taxonomy. B) One dimensional scatterplot displays the environments GROWdb MAGs were detected in across 266,764 metagenomes in the Sequence Read Archive with each point representing a single MAG. Environments are ordered by number of metagenomes GROWdb MAGs were detected in from left to right, with the number of metagenomes also noted along the x-axis. Freshwater related environments are highlighted in blue.

The dereplicated MAG database (*n* = 2,093) contained genomes from 27 phyla, many of which represent the most abundant and cosmopolitan lineages in rivers^18, 26–28^. Beyond providing new genomic resources for these ecologically “known” taxa, the GROWdb MAGs provide new genomic resources for many less well-known taxa. A subset of our genomes represented newly sampled lineages, including 10 families and 128 genera across 16 phyla (**Extended Data Fig. 2**). Additionally, a large proportion of MAGs belonged to lineages defined only by alphanumeric names (e.g., Uncultured Bacterial and Archaeal genomes, UBA^29^) at the phylum (n=1), class (n=17), order (n=121), and family (n=196) levels (**Extended Data Fig. 2**). Highlighting the relevance of GROWdb, analysis of 266,764 public metagenome datasets in the Sequence Read Archive (SRA)^25^ revealed that species in GROWdb were detected in 90% of metagenomes classified as riverine and 46% of metagenomes classified as freshwater, aquatic, or riverine (based on matching taxonomic assignments). We verified the most prevalent phyla and genera in GROWdb had parallel representation in publicly available metagenomes (**Extended Data Fig. 2**). Moreover, GROWdb species were detected from other environments including wastewater, lake water, sediment, marine, estuary, activated sludges, and soil, supporting the notion that rivers harbor diverse communities across habitats acting as integrators across landscapes (**Fig. 2B**).

The comparison to publicly available metagenomes underscored the need for this river-based microbiome study, as freshwater related metagenomes were only half to a third as many as their soil and ocean counterparts respectively in the SRA^25^. Additionally, this analysis highlighted the importance of standardized metadata practices for data reuse, as more than 10% of metagenomes in the publicly available set had vague classifications such as ‘metagenome’ or ‘bacterium’, making the data practically unusable. GROWdb ascribes to standardized protocols and metadata practices^2, 30, 31^, making interoperability a hallmark of this resource and permitting meta-analysis with other studies, which is of utmost importance as our ability to scale multi-omics methods rapidly increases.

### Aerobic and light-driven energy metabolisms are core and dominant across river microbiomes

We identified core and dominant features of riverine metagenomes and metatranscriptomes across rivers (**Fig. 3**). In terms of metagenome dynamics, members of the Actinobacteria, Proteobacteria, Bacteroidota, and Verrucomicrobiota dominated all samples (**Fig. 3A**). Within these phyla, genera that were the most cosmopolitan (occupancy; number of samples) across samples, were also the most abundant members of these communities (**Fig. 3B**). This was especially true for MAGs affiliated with the genus *Planktophilia*, a well-known freshwater microorganism^32^, that were present in 70% of the GROW metagenomes, and had the highest mean relative abundance across samples at 12%. Five other genera including *Limnohabitans A*, *Polynucleobacter*, *Methylopumilus*, *Nanopelagicus*, and *Sediminibacterium* were also present in >50% of metagenomes.

**Figure 3:**
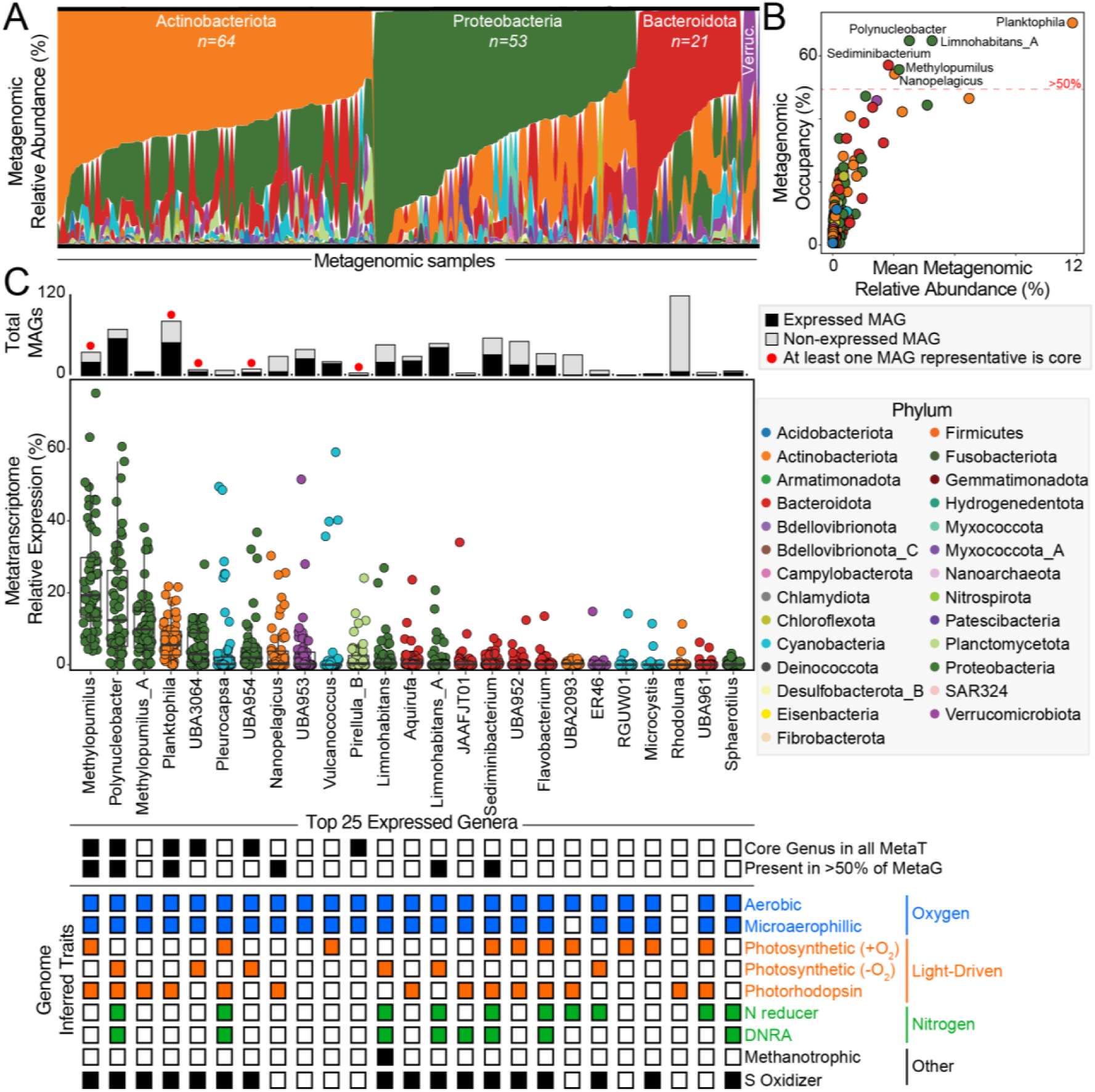
Core lineages and functions across river microbiomes. A) Phyla metagenomic relative abundance across samples, with each sample organized by the most dominant phyla from top to bottom along y-axis. Samples grouped by dominant phyla along the x-axis. Actinobacteriota, Proteobacteria, Bacteroidota, and Verrucomicrobiota (Verruc.) phyla are the most dominant across samples. B) Scatter plot highlights metagenomic relative abundance versus metagenomic occupancy (percent of metagenomes a genera was present in), with points representing each genera in GROWdb and colored by Phylum. Genera detected in more than 50% of samples (red dashed line) are named. C) The top 25 most transcribed (highest metatranscriptomic expression) genera are shown by boxplots, with each point representing a single metatranscriptome. Stacked bar chart above boxplots indicates the number of MAGs in GROWdb within each genus and colored by detection in metatranscriptomes (black, expressed; grey, non-expressed). A red circle above bar indicates that one of the genomes was core across metatranscriptomes as defined as having gene expression in every sample. For each of the top 25 expressed genera, black boxes represent those that were detected in 100% of metatranscriptomes (core genera) and >50% of metagenomes. Below, inferred genomic potential of each genera is indicated, including aerobic respiration (blue), light-driven energy metabolism (orange), nitrogen metabolism (green), and other metabolisms (e.g., methanotrophy and sulfur oxidation, black).

For the subset of samples with paired metranscriptomes, we evaluated the microorganisms that were most transcriptionally active. To focus on the most relevant lineages, we limited our analyses to MAGs that were expressing genes in at least 10% of the samples. These resulted in a quarter of the 2,093 MAGs being considered active, including at least one representee from 19 of the 27 phyla displayed in **Fig. 2**. The six most pertinent genera identified by metagenomics (**Fig. 3B**) also belonged to the top 25 genera with the highest mean gene expression (**Fig. 3C**), indicating prevalence, dominance, and activity were in agreement. Furthermore, three of these pertinent lineages (*Methylopumilus*, *Polynucleobacter, Planktophilia),* as well as members of Pirellula B, and two alphanumeric genera of Burkholderiaceae (UBA3064, UBA954) were transcriptionally active in every metatranscriptome, here denoted as the core, active genera. Notably, this was not an aggregate genus level effect, as each of these genera apart from *Polynucleobacter* had a single MAG representative that was expressed in every metatranscriptome, indicating that some microbial strains have widespread metabolic activity across rivers. Here we show how analyses of GROWdb allows us to constrain the thousands of microbial genomes to a most-wanted list of the most active microorganisms in river systems, identifying lineages and metabolic pathways that could represent diagnostic or metabolism targets needing accurate representation in biogeochemical models moving forward.

To understand the impacts of these core, transcriptionally active genera in modulating river geochemistry, we used genomic content to assign metabolic traits to each MAG, inventorying the capacity to use oxygen, light, nitrogen, sulfur and other key energy generation systems (**Extended Data Fig. 3, Extended Data File 3**). We found that the core and most expressed genera had the capacity for aerobic respiration and the use of light as an energy source, capturing energy via high-yield oxygenic or anoxygenic photosystems or simple, low yield photorhodopsins. In fact, of the top 25 most active genera, more than 90% were capable of aerobic respiration or light driven metabolisms, with many encoding multiple light harvesting mechanisms (**Fig. 3C, Extended Data Fig. 4**). In addition to heterotrophy and autotrophy, many of these core active lineages had the capacity for aerobically oxidizing inorganic electron donors like sulfur and possibly methane, the latter via a divergent particulate methane monooxygenase (**Extended Data Fig. 5**). Lastly, half of these most active genera contained the capacity for nitrogen reduction via respiration or by dissimilatory nitrate reduction to ammonium (DNRA) (**Extended Data Fig. 6**). Together the encoding of both aerobic and anaerobic energy systems, and diverse electron donors across the many core, active taxa highlight the metabolic versatility harbored in river surface waters.

Some critical river biogeochemical processes such as nitrification were represented by GROWdb MAGs but were not sampled in the top 25 most active genera. In surface waters, nitrification appeared to be catalyzed by bacteria, a finding consistent with taxonomy profiles from our unassembled reads where archaea were less than 3% of the relative abundance across samples (**Extended Data Fig. 7**). We identified one MAG within the bacterial *Nitrosomonas* genus that encoded genes for ammonia oxidation (the first step in nitrification). We note this genome also included genes to produce the greenhouse gas nitrous oxide (N_2_O), a finding consistent with other ammonia oxidizing bacteria^33^.

Two other GROWdb MAGs contained genes for nitrite oxidation (the second step in nitrification) with taxonomy assignments to the *Nitrospira_D* genus and an unassigned species within the Palsa1315 genus of the Nitrospiraceae family. With these genomes being up to 95% complete, we infer comamomox^16, 34^ is unlikely as these MAGs contained genes for nitrite oxidation but lacked genes for ammonia oxidation. These two nitrite oxidizers were detected in 14-88% of the metatranscriptome samples, including detection of transcripts for the key protein in nitrite oxidation. Each of the three nitrifier MAGs contained genes for combating reactive oxygen species (superoxide dismutase, catalase, and/or peroxidase) and a photolyase gene involved in the repair of damage caused by exposure to ultraviolet light, all adaptations likely important in surface waters^16^. Overall, our findings uncover the nitrifier metabolic potential and expression in rivers, which are underrepresented in genomic databases compared to nitrifiers from soil and marine habitats.

While not core members, we also detected 17 Patescibacterial MAGs that were transcriptionally active from the 48 total MAGs sampled in this phylum. These genomes all lacked the capacity for aerobic or anaerobic respiration and were inferred to be anoxic, obligate fermenters, consistent with prior genomic reports from this phylum that to date lacks any pure culture, characterized representatives^35, 36^. Given surface waters are oxic, we verified the abundance patterns reported here were consistent with other river metagenome and amplicon-based studies^37, 38^, where these lineages accounted for up to 7% of the relative abundance in river surface water communities. It is possible that these obligately anaerobic members exist as endosymbionts, or thrive in lower oxygen niches associated with suspended particles, biofilms or hyporheic environments where oxygen can be depleted during dissolved organic matter decomposition^39, 40^. In support of the latter, we observed that relative abundance and expression of Patescibacteria significantly decreased with river size (**Extended Data Fig. 8**), suggesting these obligate fermenters were more active in shallow waters when there is greater exchange between water and the stream bed^41^.

The occurrence of antibiotics and antibiotic resistance genes (ARGs) in riverine systems has become a growing concern worldwide. As rivers flow with heavy antibiotic burdens, antibiotic resistance develops rapidly and disseminates into various environmental compartments^42^. Antibiotic production is also part of natural competition in these complex communities. We cataloged 1,587 candidate ARGs recovered from 1135 (54.3%) MAGs in GROWdb, representing 25 different Phyla. Approximately 29% of the ARGs had evidence of expression in the metatranscriptome of one or more samples, and eleven samples had at least 20 ARGs with evidence of expression. Because our analysis was MAG-focused, these numbers may represent a floor on ARG prevalence in rivers, as they do not include plasmid-encoded ARGs. These candidate ARGs represent 25 broad antimicrobial resistance gene families as defined by the Comprehensive Antibiotic Resistance Database (CARD)^43^. Individual MAGs sometimes coded for ARGs from multiple gene families and targeting multiple drug classes.

Interestingly, most (n=1,219) candidate ARGs were homologs of proteins coded in glycopeptide resistance (*van*) gene clusters, which occurred in 955 distinct MAGs. Although glycopeptide antibiotics and self-protective resistance genes are often associated with Gram-positive actinomycete producers^44, 45^, in the GROW data 65% of the MAGs containing *van* homologs were found in 23 other diverse non-Actinobacteriota phyla, including 140 MAGs with at least 2 unique *van* gene types. Given self-protective glycopeptide resistance gene clusters have been found adjacent to or as part of a variety of different antibiotic biosynthetic gene clusters in actinobacteria^44, 45^, this diversity suggests that rivers, like soils, might harbor both unique antibiotic biosynthetic gene clusters and antibiotic-resistance genes.

### River microbiomes exhibit biogeographical patterns at the continental scale

One of the strengths of our experimental design was the spatial, chemical, and physical variables that accompanied our microbiome sampling, allowing us to contextualize the factors driving microbial biogeography at the continental scale. Previous studies have done this at local or regional levels using taxonomy alone^17, 20, 46^, but to our knowledge these analyses have not incorporated functional gene-trait information nor been done at the continental scale. We hypothesized that river microbial communities exhibit biogeographical patterns at the continental scale of inquiry, and that these patterns would be predictable from hydrobiogeochemical, geographic, and land management factors. Every sample had a paired suite of more than 250 physical, chemical, and spatial variables (e.g., stream size, latitude, total nitrogen), which we used to identify the potential drivers of microbiome structure and expressed function (**Extended Data File 1**).

Of all the river site variables examined (**Fig. 4A**), stream order, a numerical ranking of the relative river size that spans small headwater streams (low order 1-3) to larger rivers like sections of Mississippi river (high order 8-12), was the most important controller of microbiome composition. River size was more important than latitudinal position or total carbon, which are often cited as controllers of microbiomes across other habitats^47, 48^. Both metagenomes and metatranscriptomes were structured by stream order (**Fig. 4A and 4B**), providing evidence in favor of the River Continuum Concept^1^, detailed below. After stream order, expressed microbial functional profiles were also influenced by watershed air temperature (both mean and maximum), area, and total runoff (**Fig. 4A**).

**Figure 4:**
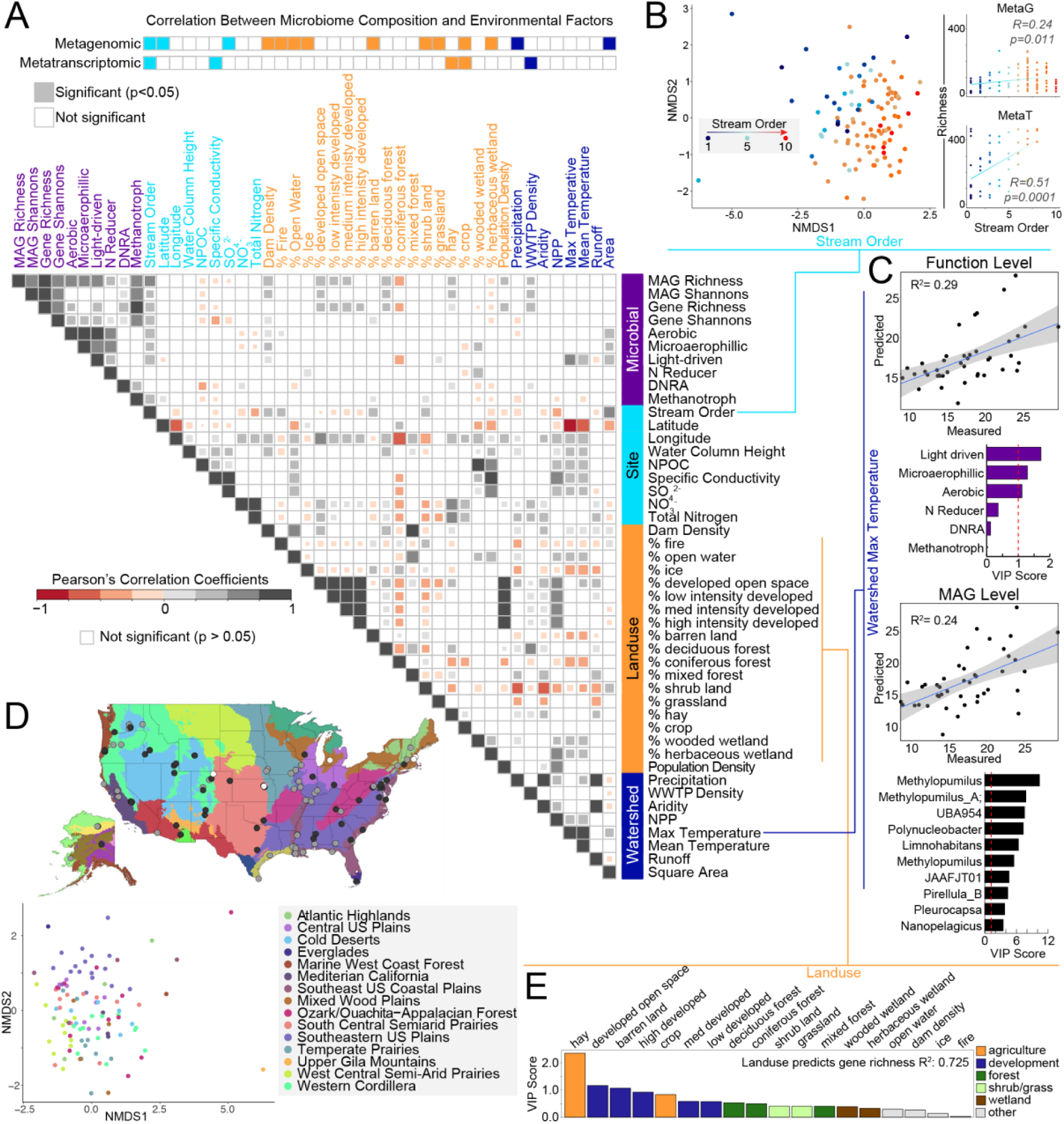
Patterns and drivers of river microbiome composition and function. A) Metagenomic and metatranscriptomic composition, function, and diversity were related to 36 selected site, land use, or watershed variables using Mantel tests (top, two rows). Below this was followed with pairwise comparisons using Pearson’s correlation (heatmap in A). Variables are colored by category including microbial (purple), site or local (light blue), land use (orange) and watershed metrics (dark blue). For pairwise comparisons of microbial data, metatranscriptomic metrics were used for diversity and function abundance calculations. B) Microbial community diversity was significantly associated with stream order as depicted by non-metric multidimensional scaling of genome resolved metagenomic Bray-Curtis distances (left, beta-diversity) and Pearson correlations of richness to stream order (right, alpha-diversity) with points colored by stream order. C) Sparse Partial Least Squares (sPLS) regressions show significant function (top) and MAG level (bottom) expression predictions of watershed maximum temperature, with key variables (Variable Importance Projection >1) denoted in bar graphs below. D) Non-metric multidimensional scaling of genome resolved metagenomic Bray-Curtis distances shows clustering of microbial communities by ecoregion (classified by Omernik II), with sampling location depicted on map above (mrpp, p<0.001). E) Random forest analysis showed that land use metrics predicted expressed gene richness with 72% accuracy, with order of variable importance shown as a bar graph. Abbreviations: NPOC, Non-Purgable Organic Carbon; DNRA, Dissimilatory Nitrite Reduction to Ammonia; WWTP Density, Waste Water Treatment Plant Density; NPP, Net Primary Production.

Given this relationship with local atmospheric conditions, we sought to understand which functional traits and microorganisms most contributed to these community level observations. Regression based modeling showed light driven metabolisms, followed by aerobic processes, were the most important variables, predictive of mean and max watershed air temperature (**Fig. 4C**). The most important organismal predictors of maximum watershed temperature were the core active lineages like *Methylopumilis*, UBA954, *Polynucleobacter*, and *Limnohabitans* that were actively transcribing genes for light harvesting metabolisms **(Fig. 3C)**. Our findings show that light harvesting metabolisms are critical to energy generation in rivers and suggest that climate influences on water temperature play a defining role in the niches of these microorganisms. These findings are consistent with reports from marine systems^49^, hinting at an emerging rule set shared across aquatic microbiomes.

Beyond environmental factors we also observed that geographic position played a role in structuring river microbiomes. For instance, microbial community genomic membership was structured across ecoregions defined by Omernik level II classifications^50^, with drier climate mixed grass influenced river microbiomes sharing similar microbial communities that were distinct from those derived from wet to sub-tropical regions (**Fig. 4D**). Similarly, hydrologic unit code (HUC), a classification system for watersheds in the United States shown in **Fig. 1C**, recognized distinct microbial communities from continental subregions (**Extended Data Fig. 9**). These findings support earlier work examining stream microbial succession, which showed that river microbial communities are inoculated from the landscape, and this terrestrial influence continues to play an important role in downstream community assembly processes^19^. We note that the spatial structuring was not observed at the expressed functional level, indicating species changes are compensated by functional equivalence at this continental scale. This finding suggests taxonomic information may not be best suited for translation of microbiome content into management indicators, unless incorporated into an eco-regional framework as has been suggested for soil health indicators^51, 52^.

To use microbiota information as sentinels for monitoring human and environmental health in river systems, a greater understanding of bacterial community structure, function, and variability in lotic systems is required^53^. Particularly important is how different land management types may structure surface water microbiomes. Here we used a machine learning approach to associate river MAGs or gene content and expression with different land management regimes. We found that land use predicted metatranscriptomics richness (72% accuracy), with agriculture and urban influences being the most predictive variables (**Fig. 4E**). Together our findings show how river microbiomes are driven by multiple environmental factors, including local, watershed, regional, and continental factors. Analyses using GROWdb provide a new framework for the environmental factors and determinant mechanisms that shape riverine communities.

### Microbial gene expression provides evidence for the river continuum concept

The River Continuum Concept (RCC) provides a framework for integrating predictable and observable biological features of flowing water systems, and further characterizing how biodiversity changes along a river system^1^. Specifically, the RCC postulates that as rivers increase in size, the influences of terrestrial inputs will decrease. It also assumes, that biological richness will initially increase with stream order complexity due to maximum interface with the landscape, but then decrease along with river width and discharge. Support for the applicability of the RCC to microbial communities has been observed as decreased microbial 16S rRNA gene richness occurring across stream order gradients in Toolik^19^, Danube^54^, Mississippi^53^, and Amazon^55^ rivers. Here we show that the RCC also applies to microbial community genomic and transcriptomic patterns at the continental scale (**Fig. 4B**). Importantly, this discovery provides an opportunity to interrogate diversity and functional trait dimensions along a river continuum at broader scales.

First, we were interested in how microbial richness at the metagenome and metatranscriptome level changed across the stream order gradient, and if these followed rules like 16S rRNA studies from single rivers. At the metagenome level, overall genome richness had a positive correlation with stream order, peaking at a stream order six (**Fig. 4B**). At the metatranscriptome level, richness had a strong positive correlation with stream order and peaked at stream order 8, the highest stream order profiled by metatranscriptomics (**Fig. 4B**). Metagenome results were consistent with previous reports of the RCC where stream order peaks in mid-sized streams^53^. To our knowledge this is the first report of genome-resolved metatranscriptomics across rivers and suggests that genome inferred transcriptional richness may be governed by a different set of environmental controls than gene presence at the continental scale.

One major control on biological diversity described by the RCC is variability in sunlight exposure. Lower order streams are often characterized by thick shore vegetation or overhanging trees that limit sunlight penetration and restrict phytoplankton and benthic microalgae primary production^1, 56^. Consistent with this idea, we observed statistically significant increase in light-driven microbial metabolisms when moving from lower order streams to higher order rivers (**Fig. 4A**). Additionally, the RCC proposes the ratio of photosynthesis to respiration (P/R) increases in medium-larger rivers but is decreased in the smallest and largest rivers due to light limitations from riparian vegetation occlusion and turbidity, respectively. Using microbial gene expression coupled to genome-resolved lifestyle information we estimated P/R ratios, revealing the highest P/R ratio in rivers with stream orders of 6-8, providing tentative support for this concept. However, the robustness of this P/R indicator would need further evaluation in larger ordered rivers (e.g., 9-12) rivers, which are under sampled in this metatranscriptome dataset.

Another ecological control described by the RCC is a downstream decrease in the importance of terrestrial carbon inputs. We hypothesized gene expression would show that microbial carbon usage reflects decreasing impacts of terrestrial inputs with river size. To resolve changes in microbial metabolism across a stream order gradient, we defined carbon usage patterns based on microbial gene expression in GROWdb MAGs. Our findings show significant differences in expressed microbial carbon usage following stream order gradient (**Fig. 5A**). Specifically, transcript of genes targeting polymers, aromatics, and sugars are upregulated in low order streams, while methylotrophy gene transcripts, primarily from methanol oxidation, increased in higher order rivers. We consider methanol is likely autochthonous, derived from river phytoplankton biomass^57^ or microbial metabolism of aromatic allochthonous plant litter^58, 59^. Our findings show that the inferred microbial metabolisms related to carbon usage follow the expected decrease in impact of terrestrial inputs proposed by the RCC, but we acknowledge more research is needed to validate these new insights, especially from higher order rivers.

**Figure 5:**
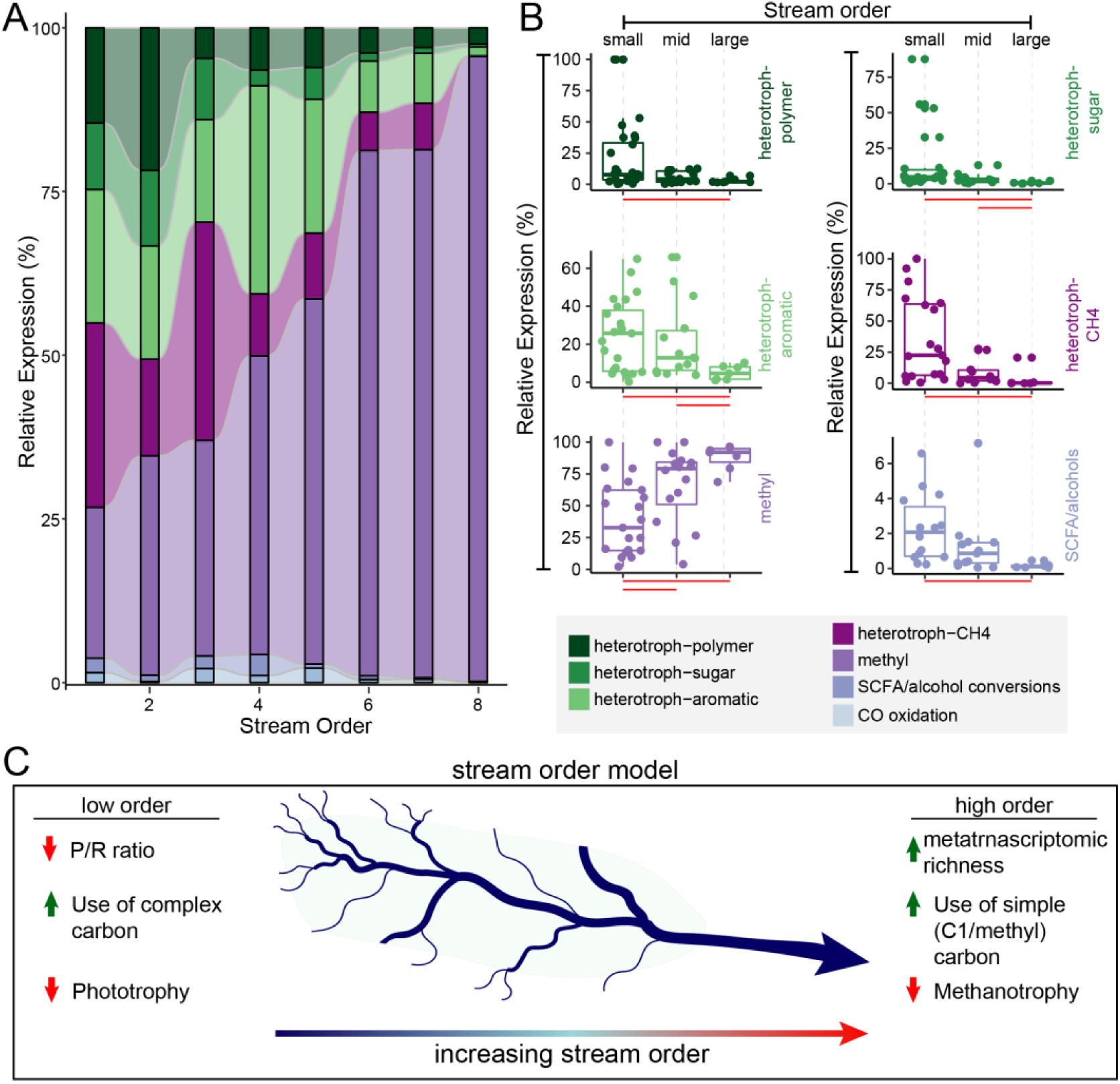
Microbial lifestyle and carbon utilization are structured along a stream order gradient. A) Alluvial plot shows the relative expression of microbial lifestyles (defined in methods) across stream order gradient. B) One dimensional scatterplots correspond to data in A, with each point representing a single sample and streams grouped by small (1-3), mid (4-6), and large (7-8) orders. Significant differences in expression (p<0.01) between small, mid, and large order streams are denoted by horizontal red bars below each plot. C) The stream order model highlights changes in microbial expression from small (left) to large (right) order streams.

In summary, river systems were once thought of as passive pipes, transporting water from terrestrial to marine systems. As a result, it was regarded that microbiomes in these systems were a mixture of randomly assembled microorganisms with little predictive capability. Instead, we show river microbiomes and encoded functionalities are not haphazardly distributed but are instead structured by river size, ecological region, and land management regimes. This study also supports the application of the RCC to microbial communities and provides the first evidence that landscape patterns in river microbiomes are grounded in mechanistic changes in genomic function. We show that microbial richness both in terms of genome potential and expression, as well as expressed functional attributes, follow RCC tenets and are molded by the physical-geomorphic environment. This application of GROWdb to the RCC adds a new view of how microbial metabolism changes across rivers.

### Conclusion

Rivers are a critical part of the Earth System for their roles in dispersing water across terrestrial ecosystems. Changing climate impacts rivers via altered precipitation intensity, surface runoff, flooding, fires, sea level rise, and droughts, and all of these have direct impacts to human health, agriculture, energy production, and ecosystem resiliency^60^. Additionally, 2/3 of drinking water in the United States come from surface river waters. Consequently, river management is expected to be one of the most politically charged topics in decades to come^61^. Microbes are master orchestrators of nutrient and energy flows that will likely dictate water quality under current and future water scenarios.

The Earth System is at a breaking point and microbiomes may be one of the levers to push back on climate chaos^4^. The field of environmental microbiology has undergone rapid transition from microscopy revealing abundant microbes to gene markers demonstrating the vast diversity of microbiota colonizing Earth’s habitats^62^. Today’s challenge is linking “who is there” to their functional and ecosystem roles and doing so at scale that allows elucidation of emergent properties from diverse and complex systems. Here we crowdsourced a large-scale river microbiome study using standardized sampling, processing, sequencing, and analysis to enable cross-site comparisons and modular augmentation. This resulted in a comprehensive resource that includes a collection of over two thousand curated metagenome-assembled genomes. We show this resource can be used to rapidly contextualize “who” and “what processes” microbes are modulating in river systems. This product and its many data access and synthesis sites reduces the computational barriers for expediting the translation of reads to functional content. Ultimately, GROWdb is a highly contextualized, genome-resolved resource spanning river sizes, locations, and land management regimes to promote microbiota derived insights across disciplinary boundaries.

GROWdb offers a genome-centric window into river microbiota and a FAIR-use cyberinfrastructure-powered platform for future researchers. GROWdb analyses presented here reveals core functionalities of river microbiomes and illuminates key metabolisms and microorganisms that are conserved and are transcriptionally active across the United States river systems. We envision that this genomic infrastructure will pave the way for future developments in water quality monitoring and identifying biomarkers indicative of land use or water quality changes. Collectively, GROWdb fills a major knowledge gap in the current understanding of microbial diversity and function in river ecosystems, observations that can integrated into predictive watershed scale models.

## Methods

### Sample collection through crowdsourcing and standardization of workflows

To build GROWdb, we used two approaches to obtain samples from across United States rivers. One was a network-of networks^21^ approach based on sampling efforts of the Worldwide Hydrobiogeochemistry Observation Network for Dynamic River Systems (WHONDRS) consortium^63^, which is designed to facilitate the development of transferable scientific understanding and mutual benefit across stakeholders^30, 31^. The WHONDRS sampling itself was based on sending free sampling kits, along with standardized protocols, to interested researchers globally (sites outside of North America are not included in GROWdb). These researchers volunteered their time to collect samples and sent the samples back for processing using consistent methods to enable cross-site comparisons, interoperable data, and transferable understanding. Samples from the WHONDRS consortium contributed 44% of the metagenomes and all the metatranscriptomes in GROWdb. The other approach was through a collaboration with the U.S. Geological Survey (USGS) National Water Quality Network (NWQN)^64^. The rest of the samples were collected under the NWQN campaign, which is a USGS long-term water-quality monitoring program to characterize consistent information on streamflow and water-quality conditions. Data are collected to assess the status and trends of water-quality conditions at large inland and coastal river sites, as well as in small streams indicative of urban, agricultural, and reference conditions^64^.

Samples described in GROWdb version 1 were collected under the WHONDRS 2019 sampling campaign (**Extended Data File 1**) and described in Garayburu-Caruso, et al.^65^ Briefly, at each sampling location, collaborators selected sampling sites within 100 m of a station that measured river discharge, height, or pressure. Geochemical data collected under the WHONDRS 2019 sampling campaign are available ESS-DIVE, with methods described^24^. For microbiome analyses, at each site, approximately 1L of surface water was sampled using a 60 mL syringe and was filtered through a 0.22 μm sterivex filter (EMD Millipore). Filters were capped, filled with 3mL of RNAlater and shipped to Pacific Northwest National Laboratory on blue ice within 24 hours of collection. Surface water samples and filters were immediately frozen at −20 °C upon receiving for nucleic acid extraction, respectively. Methods of sample collection used by the National Water Quality Network (NWQN) conform to the USGS National Field Manual for the Collection of Water-Quality Data^66^, and DNA was collected on 0.22 μm Sterivex-GP filter (EMD Millipore). Here we provided kits integrated with USGS protocols for river sample processing with samples were preserved as described previously^67^. All samples were stored on ice and stored at –20°C until nucleic acid extraction.

A key component of this analysis was the standardization that occurred in data processing and analyses. DNA and RNA were co-extracted at single facility at Colorado State University. DNA and RNA were coextracted from filters at Colorado State University using ZymoBIOMICS DNA/RNA Miniprep Kit (Zymo Research Cat. # R2002) coupled with RNA Clean & Concentrator-5 (Zymo Research Cat. # R1013). Samples were eluted in 40 μL and stored at −20 °C until sequencing. Similarly, a Community Sequencing Project (CSP) provided by the Joint Genome Institute (JGI) ensured that sequencing protocols and methodologies were consistent across the project. All the metagenomes and 23% of the metatranascriptomes were provided by JGI, with the balance of metatranscriptomes processed at University of Colorado Anschutz using the same kits and methods as specified by JGI. Lastly, sequence data processing for each sample was performed using identical methods, with the GROWdb standard operating procedures documented on GitHub^68^. Collectively, the use of crowdsourced approaches, JGI support, and standardized methodologies resulted in GROWdb, a compendium of river microbiome data, an endeavor that would not have been possible to execute in this time frame by a single laboratory alone.

### Acquisition of geospatial data

The watershed statistics for each sample were primarily obtained from the Environmental Protection Agency’s StreamCat database^69^ and the National Hydrography Plus Version 2 (NHDPlus V2) Dataset using the *nhdplusTool*s package^70^ in R. StreamCat provides over 600 consistently computed watershed metrics for all waterbodies identified in the US Geological Survey (USGS)’s NHDPlusV2 geospatial framework, making it a suitable data source for the broad spectrum of sample locations in this study. For watershed metrics that were not included in StreamCat (i.e., dominant Omernik Ecoregion, mean net primary production, and mean aridity index), we first delineated each sample’s watershed using *nhdplusTools*, then utilized the *terra* package^71^ to aggregate the additional datasets across each site’s watershed accordingly. This approach is consistent with SteamCat’s geospatial methodology.

Lastly, we collected streamflow data for sites that had a nearby stream gauge. For locations without an identified co-located stream gauge (WHONDRS typically co-located their sample sites with a stream gage), we identified USGS stream gauges within 10 kilometers up- or downstream of our sampling locations using the *dataRetrieval* and *nhdplusTools* packages. All stream gages were then manually verified for their applicability to each sampling site (e.g., verifying there were no dams between the site and the stream gage, a major confluence, etc.). See **Extended Data File 1** for a complete list of datasets included in our analysis. The complete R workflow for this geospatial analysis can be found on GitHub^72^.

### Metagenomic assembly, binning, and annotation

At the Joint Genome Institute, genomic DNA was prepared for metagenomic sequencing using Plate-based DNA library preparation on the PerkinElmer Sciclone NGS robotic liquid handling system. Briefly, one nanogram of DNA was fragmented and adapter ligated using the Nextera XT kit (Illumina) and unique 8bp dual-index adapters (IDT, custom design). The ligated DNA fragments were enriched with 12 cycles of PCR and purified using Coastal Genomics Ranger high throughput agarose gel electrophoresis size selection to 450-600bp. The prepared libraries were sequenced using Illumina NovaSeq sequencer following a 2×150nt indexed run recipe.

Resulting fastq files were assembled and binned using the accessible GROWdb pipelines released on GitHub^68^. Briefly, to maximize genome recovery three assemblies were performed on each set of fastq files and binned separately: (1) Read trimming with sickle (v1.33)^73^, assembly with megahit (v1.2.9)^74^, and binning with metabat2^75^ (2.12.1) (2) Read trimming with sickle (v1.33)^73^, random filtering to 25% of reads, assembly with idba-ud^76^ (1.1.0), and binning with metabat2^75^ (2.12.1) (3) Bins derived from the JGI-IMG pipeline^77^ were downloaded. All resulting bins were assessed for quality using checkM^78^ (v1.1.2) and medium and high-quality MAGs with >50% completion and <10% contamination were retained. The resulting 3,284 MAGs across all samples and assemblies were dereplicated at 99% identity using dRep^79^ (v2.6.2) to obtain the dereplicated first version of the GROW database (n=2,093 MAGs). MAG taxonomy was assigned using GTDB-tk^22^ (v2.1.1, r207) and annotated using DRAM (v1.4.4)^80^.

To quantify MAG relative abundance across samples, trimmed metagenomic reads were mapped to the dereplicated MAG set using Bowtie2^81^ and output as sam files which were then converted to sorted bam files using samtools. Sorted bam files were then filtered to paired reads only with a 95% identity match using reformat.sh. To obtain the mean coverage for each MAG, we used CoverM^82^ (-m trimmed_mean). The mean coverage table was then filtered to MAGs that had at least 60% coverage across a MAG with at least 3X coverage within a sample, using additional CoverM^82^ outputs (-m relative_abundance –min-covered-fraction 0.6 and -m reads_per_base, respectively). CoverM outputs were merged in R, with script available on the GROWdb GitHub^68^.

### Metatranscriptomic mapping and analysis

RNA was prepared for metatranscriptome sequencing according to JGI established protocols. Briefly, rRNA was removed from 10 ng of total RNA using Qiagen FastSelect probe sets for bacterial, yeast, and plant rRNA depletion (Qiagen) with RNA blocking oligo technology. The fragmented and rRNA-depleted RNA was reverse transcribed to create first strand cDNA using Illumina TruSeq Stranded mRNA Library prep kit (Illumina) followed by second strand cDNA synthesis which incorporates dUTP to quench the second strand during amplification. The double stranded cDNA fragments were then A-tailed and ligated to JGI dual indexed Y-adapters, followed with an enrichment of the library by 13 cycles of PCR. The prepared libraries were quantified using KAPA Biosystems’ next-generation sequencing library qPCR kit and run on a Roche LightCycler 480 real-time PCR instrument. Sequencing of the flowcell was performed on the Illumina NovaSeq sequencer following a 2×150nt indexed run recipe.

Resulting fastq files were mapped via Bowtie2^81^ (-D 10 -R 2 -N 1 -L 22 -i S,0,2.50) to the dereplicated GROWdb. Sam files were transformed to bam files using samtools, filtered to 97% id using reformat.sh and name sorted using samtools. Transcripts were counted each gene with feature-counts^83^. Counts were transformed to geTMM (gene length corrected trimmed mean of M-values) in R using edgeR package^84^. Genes and bins were considered if they were expressed in 10% of samples. Core calculations in **Fig. 3** had an additional requirement to express at least 20 genes.

### Microbial metabolism trait and carbon usage classification

To classify microbial genes and genomes based on their carbon metabolism we curated the metabolism assignments made by DRAM^80^ using rulesets to assign genomes to functional guilds (**Extended Data Fig. 3**). For example, genomes were classified by respiratory capacity based on the presence of >50% of the subunits required for Complex 1 of the electron transport chain and the presence at least one gene for an electron acceptor. As such, for a genome to be classified as a microaerophile, we required the genome to have more than 50% of complex 1 subunit and at least one subunit of a low affinity cytochrome oxidase. Likewise, if a genome did not have more than 50% of the subunits required for Complex 1 of the electron transport chain or the potential for any electron acceptor, it was classified as an obligate fermenter (**Extended Data Fig. 3**).

From the DRAM output, we further assigned genomes as capable of carbon fixation if they encoded >70% of one of six seven carbon fixation pathways. We then assigned each MAG in each river metatranscriptome as a photoautotroph, photoheterotroph, chemolithoautotrophy, heterotroph, or mixotroph by assessing the gene expression in that system. We then focused in on genes required for utilizing different carbon substrates in the genomes identified for heterotrophy. We assigned carbon gene expression into the following categories: polymer, sugar, aromatic compound, methanotrophy, methylotrophy, short chain fatty acid utilization, and carbon monoxide utilization using DRAM assigned rules. Carbon usage curation scripts are available on the GROWdb GitHub^68^. P/R ratios were defined by the ratio of expression of light driven energy metabolisms (aerobic photosynthesis, anaerobic photosynthesis, and photorhodopsins) divided by aerobic respiration metabolisms (aerobic respiration and microaerophilic respiration).

Phylogenetic analyses were performed to refine the annotation of nitrogen related metabolism including genes annotated as respiratory nitrate reductase (*nar*), nitrite oxidoreductase (*nxr*), ammonia monooxygenase (*amo*), or methane monooxygenase (*pmo*) to improve the assignment the nitrogen cycling capabilities of GROW MAGs (**Extended Data File 3**). Specifically, Nxr/Nar and PmoA/AmoA amino acid reference sequences were downloaded^34, 85, 86^ and this set of reference sequences were combined with amino acid sequences of homologs from the GROWdb, aligned separately using MUSCLE (v3.8.31), and run through an in-house script for generating phylogenetic trees^87^. This resulted in two phylogenies, one for Nxr/Nar and one for Pmo/Amo, with this homology based approach used to refine the homology based gene annotations in the MAG database.

For *in silico* predictions of antimicrobial resistance genes (ARGs) GROWdb predicted proteins were searched for homology to proteins in the Comprehensive Antibiotic Resistance Database (CARD; v 3.2.7, downloaded June 2023) using the Resistance Gene Identifier (RGI; v 6.0.2)^43^. RGI was run locally in protein input mode with distributed input and default parameters, and with the “include loose” option. However, the final list of candidate ARGs analyzed here only includes proteins identified by RGI as “Perfect” or “Strict” hits, and only includes protein homolog models (i.e., no protein variant models were included in the analysis).

### Sequence Read Archive Analysis

To analyze the distribution of species recovered by GROW across public datasets, the Sandpiper database (https://sandpiper.qut.edu.au) was used as a basis. At the time of analysis, it contained metagenomes that were publicly available on Dec 15, 2021 (Woodcroft et. al., unpublished). Reanalysis of these datasets was carried out with SingleM 1.0.0beta7 (Woodcroft et. al., unpublished, https://github.com/wwood/singlem). The “supplement” subcommand was first used to add 95% ANI dereplicated GROW MAGs to the SingleM reference metapackage built with GTDB RS07-207 (DOI: 10.5281/zenodo.7582579). The “renew” subcommand was then used to reanalyze all metagenomes present in the Sandpiper database, outputting a taxonomic profile, detailing the species and unclassified lineages in each metagenome, together with their relative abundance.

To search for public metagenomes where GROW MAGs were present, taxonomic profiles of metagenomes containing species that had an associated GROW MAG (either novel or already represented in GTDB) were further analyzed. To reduce the incidence of false identification, we required at least 2 species represented by a GROW MAG to be present and the combined relative abundance of these species to be >1%. Metadata of metagenomes containing GROW MAGs were gathered using Kingfisher “annotate” (https://github.com/wwood/kingfisher-download).

### Statistical Analysis

All data analysis and visualization was done in R (v4.2.1) with the following packages: stats (v4.1.1), vegan (v2.6), ggplot2 (v.3.3.6), ComplexUpset (v2.8.0), tidyr (v1.2.0), dplyr (v1.0.9), corrplot (v0.92), pheatmap (v1.0.12), RColorBrewer (v1.1-3), pls (v2.8), edgeR (v3.16). Scripts for figure generation and data analysis are available on GitHub^68^.

## Data accessibility

The data underlying GROWdb are accessible across multiple platforms to ensure many levels of data use and structure are widely available. First, all reads and MAGs are publicly hosted on National Center for Biotechnology (NCBI) under Bioproject PRJNA946291. Second, all data presented in this manuscript including MAG annotations, phylogenetic tree files, antibiotic resistance gene database files, and expression data tables are available in Zenodo (https://doi.org/10.5281/zenodo.8173287).

Beyond the content listed above, our aim for GROWdb was to maximize data use by making the data available in searchable and interactive platforms including the National Microbiome Data Collaborative (NMDC)^2, 23^ data portal, the Department of Energy’s Systems Biology Knowledgebase (KBase)^3^, and a GROW specific user interface released here, GROWdb Explorer. Each platform provides different ways to interact with data in the GROWdb:

- *NMDC* GROWdb was a flagship project for the newly formed NMDC. Specifically, individual GROWdb datasets (metagenomes, metatranscriptomes, etc) are easily accessible and searchable through the NMDC data portal (https://data.microbiomedata.org/), where they are systematically connected to each other and to a rich suite of sample information, other data collected on the same samples, and standard analysis results, following Findable, Accessible, Interoperable, and Reusable (FAIR) data practices^30^.
- *KBase* GROWdb is a publicly available collection within KBase^3^, with samples, MAGs, and corresponding genome scale metabolic models found in the KBase narrative structure (https://doi.org/10.25982/109073.30/1895615). Access within KBase allows for immediate access and reuse of data, including comparison to private data analyses using KBase’s 500+ analysis tools, in a point and click format.
- *GROWdb Explorer* is a graphical user interface built through the Colorado State University Geospatial Centroid (https://geocentroid.shinyapps.io/GROWdatabase/), allowing users to search and graph microbial and spatial data simultaneously. Here the microbial data was distilled into functional gene information, so that biogeochemical contributions and the microorganisms catalyzing them can be assessed and visualized rapidly across the dataset.

In summary, GROWdb represents the first publicly available genome collection from rivers and offers data that can be leveraged across microbiome studies. GROWdb is an expanding repository to incorporate and unify global river multi-omic data for the future.

## Code availability

All scripts involved with microbial data generation, processing, curation, and visualization are available on GitHub (https://github.com/jmikayla1991/Genome-Resolved-Open-Watersheds-database-GROWdb/tree/main). Code for geospatial analysis and GROWdb Explorer are available on GitHub (https://github.com/rossyndicate/GROWdb).

## Extended Data Figures

**Extended Data Fig. 1:**
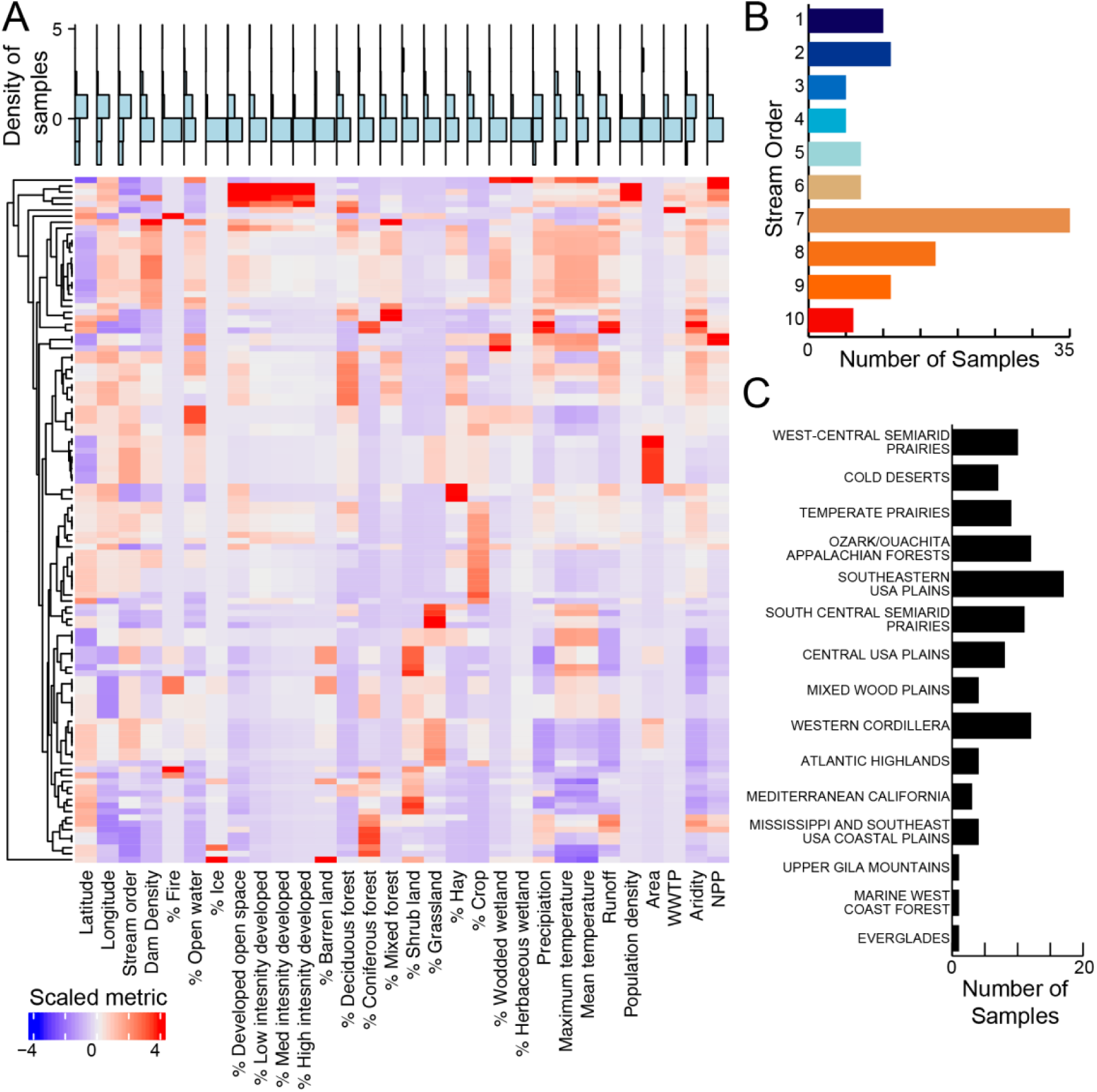
Distribution of river characteristics sampled in GROWdb. A) Heatmap of geospatial parameters sampled in GROWdb, where columns are environmental variables and rows are corresponding samples within GROWdb. Each variable has been scaled by subtracting the vector mean for each variable and dividing by its standard deviation. Blue histogram plots above highlight the distribution of samples for each variable, with high values at the top of the plot. Histogram plots of key variables used throughout the main text including stream order (B) and ecoregion (C) are also shown.

**Extended Data Fig. 2:**
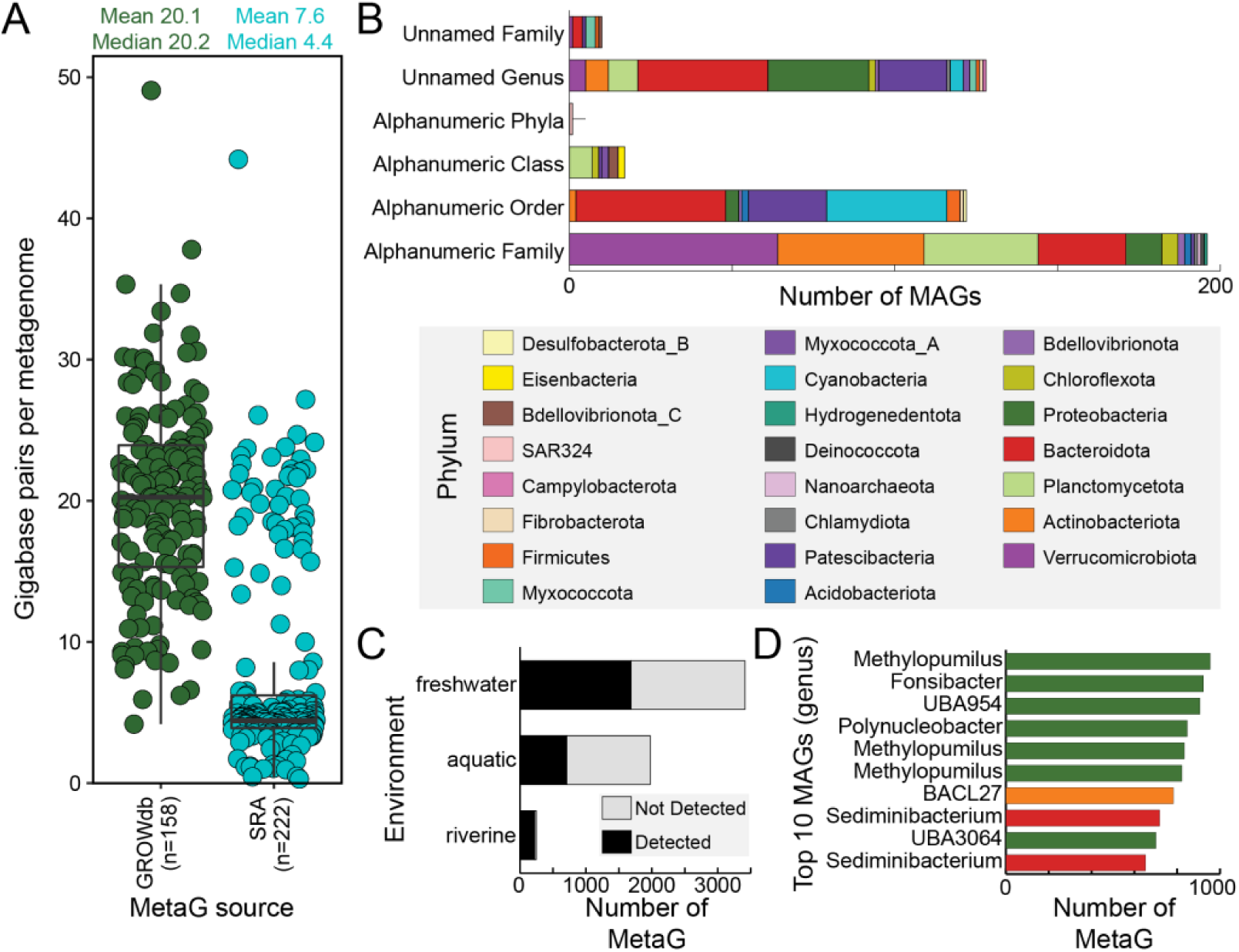
GROWdb comparison to other omics data sources. A) Sequencing depth comparison of GROWdb metagenomes (n=158) to SRA metagenomes classified as riverine (n=222) shows 3x increase in average sequencing depth for GROWdb. Each point represents a single metagenome, with mean and median values listed at the top of the graph. B) Stacked bar chart shows novelty of GROWdb MAGs when compared to GTDB (r207). Each MAG was placed at the highest level of novelty, with no assignment within a taxonomic level (e.g., unnamed family or genus) being highest level of novelty and alpha numeric identifiers being the second highest (e.g., UBA lineages). Bars are colored by Phylum. C) Stacked bar chart shows the proportion of SRA metagenomes that a GROWdb species was detected (black) or not detected (grey) within an SRA environment category. D) The top ten MAGs most frequently detected across river surface water related SRA metagenomes are displayed at the genus level on the bar chart, with colors denoting phyla (key above).

**Extended Data Fig. 3:**
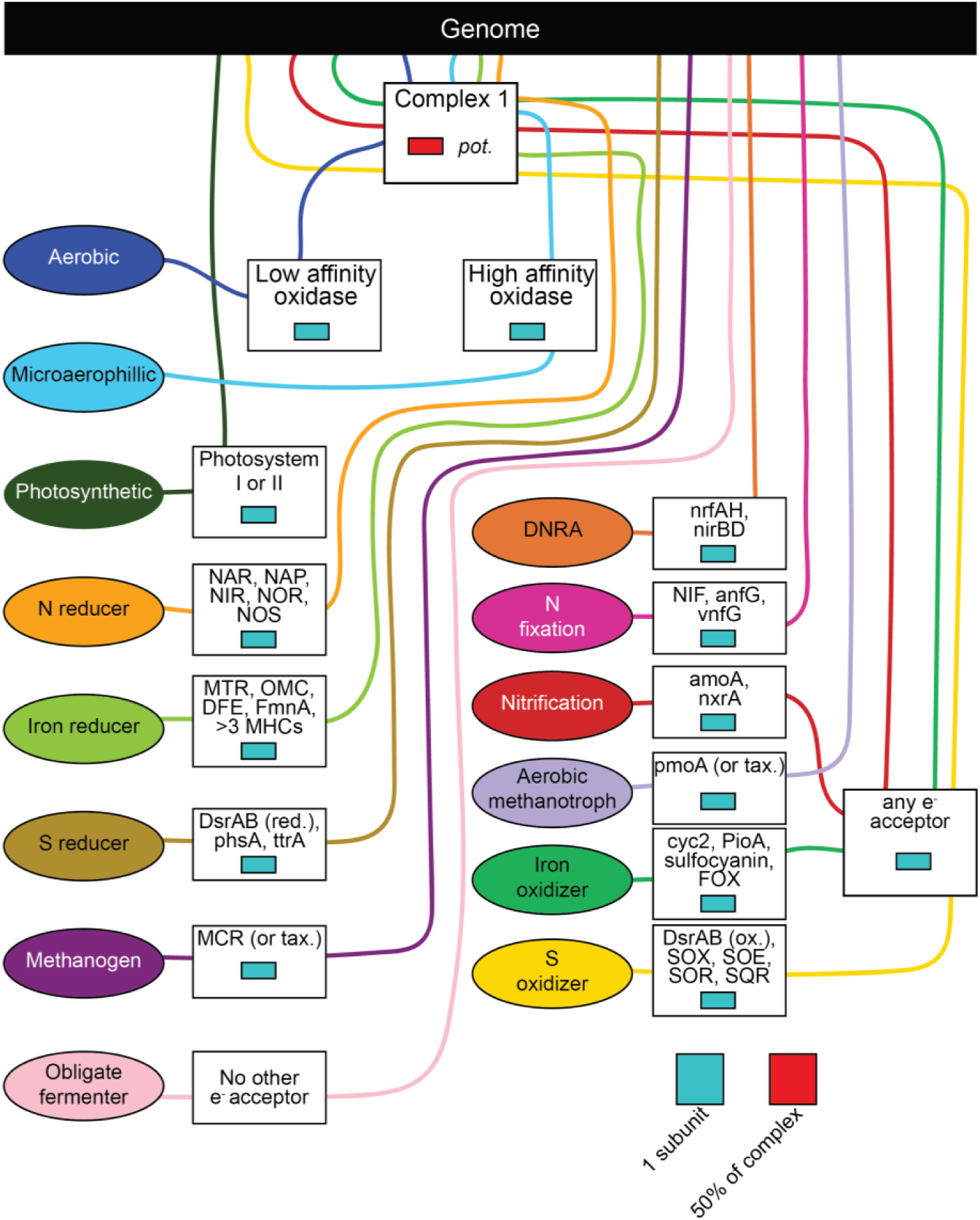
Metabolic trait assignment ruleset. Each trait is defined by a set of genes and the percent of genes required for that function. Lines flow from the genome (top black box) to traits (ovals), passing through boxes of gene requirements to be consider TRUE for that particular trait.

**Extended Data Fig. 4:**
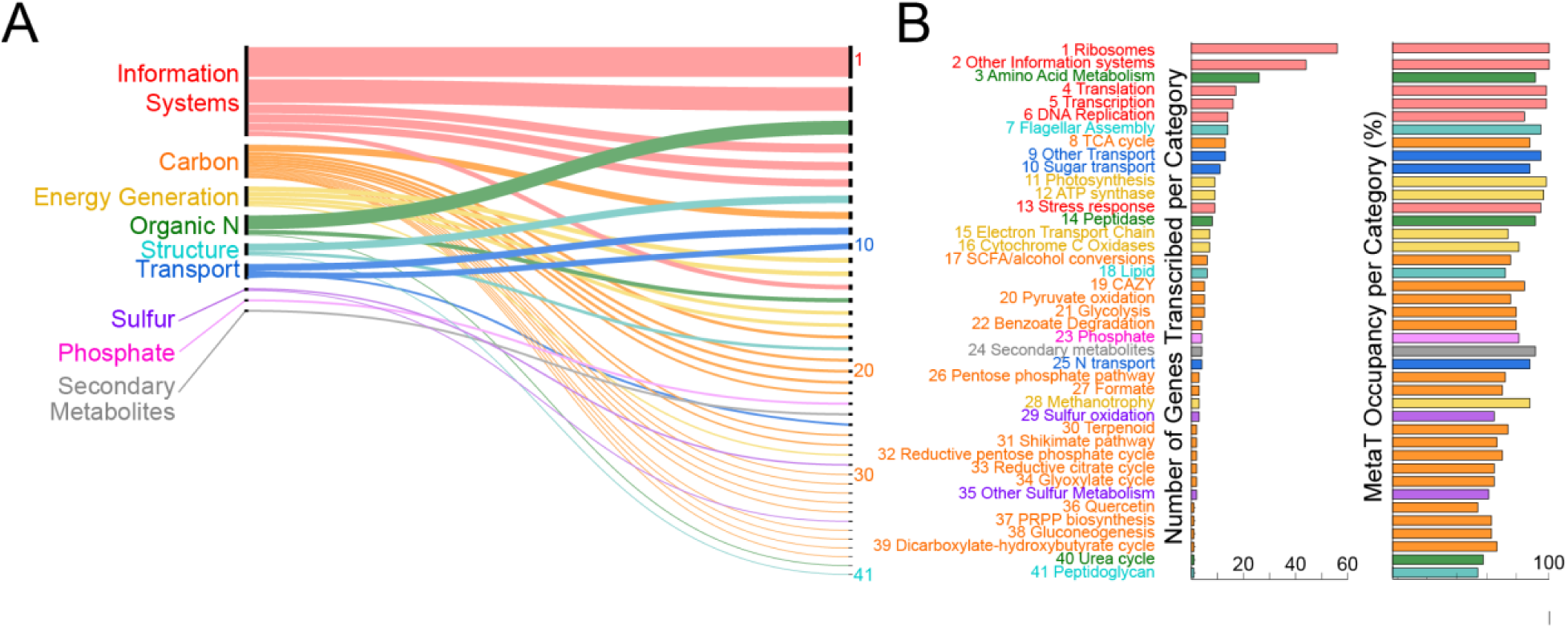
Gene level expression across rivers. Genes detected in more than 50% of metatranscriptomes, with gene functions (n=365) grouped by broad categories (n=9, A) and refined to subcategories (n=41, B). Thickness of lines and line order in A show the number of functions within a particular category (right) and subcategory (left). A and B are linked by subcategory number (1-41). For each of the 41 subcategories, the number of genes and occupancy defined as the percentage of samples detected across metatranscriptomes is shown by bar charts. Hypothetical and genes with unknown annotations are not shown, albeit 21 genes with these annotations were considered core or expressed in all metatranscriptomes.

**Extended Data Fig. 5:**
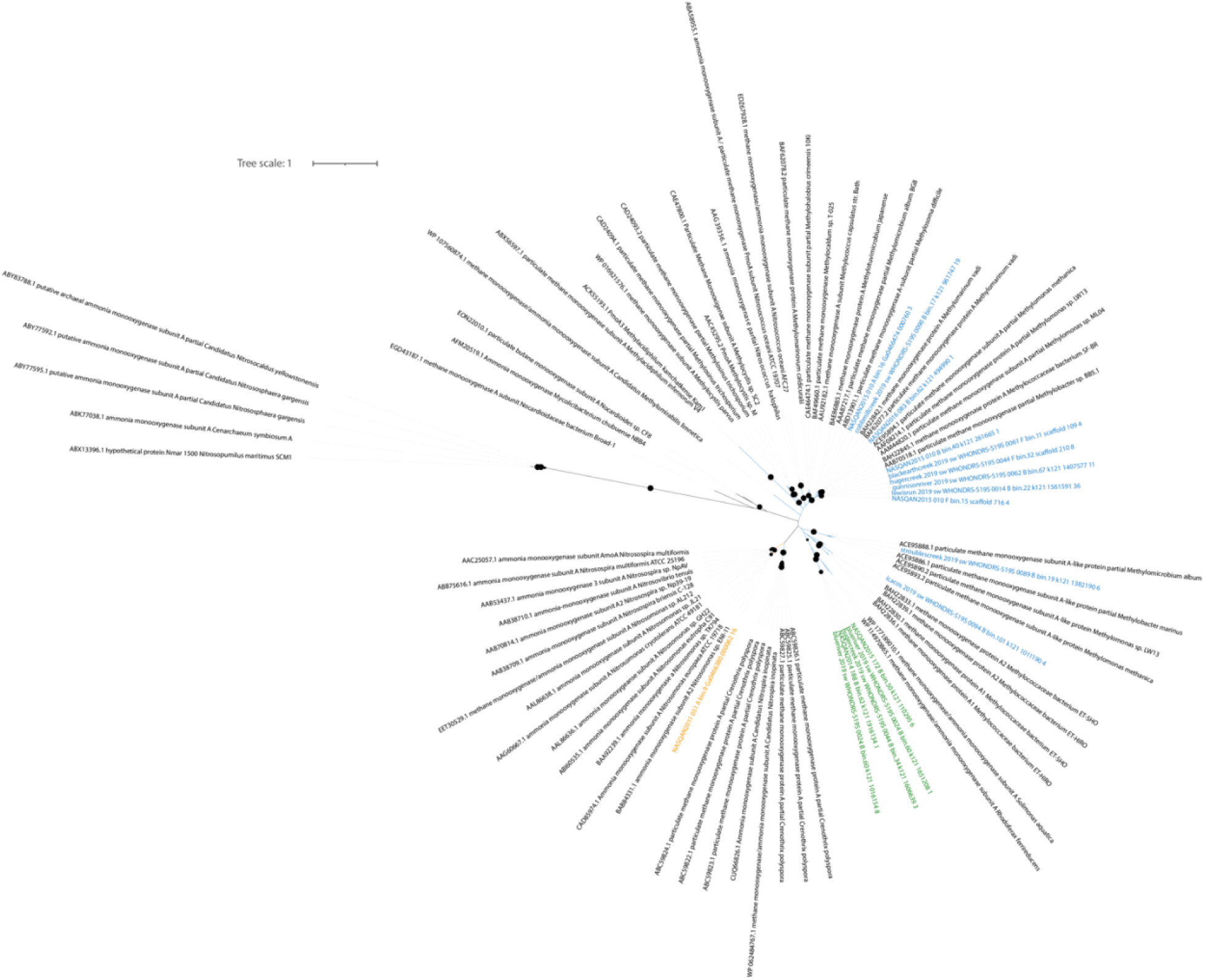
Phylogenetic analysis of ammonia monooxygenase (amoA) and methane monooxygenase (pmoA) is shown. GROWdb sequences colored by putative substrate, with ammonia (orange), methane (blue), and unknown (green), while reference sequences are shown in black text. Bootstraps >80 are shown with closed black circles at nodes. Several GROW Limnohabitans genomes contained a putative amoA/pmoA gene (similar to other *Limnohabitans* genomes), but the substrate specificity has not been confirmed, however here based on tree placement we denote these as methanotrophs. Tree files are available on Zenodo (https://doi.org/10.5281/zenodo.8173287).

**Extended Data Fig. 6:**
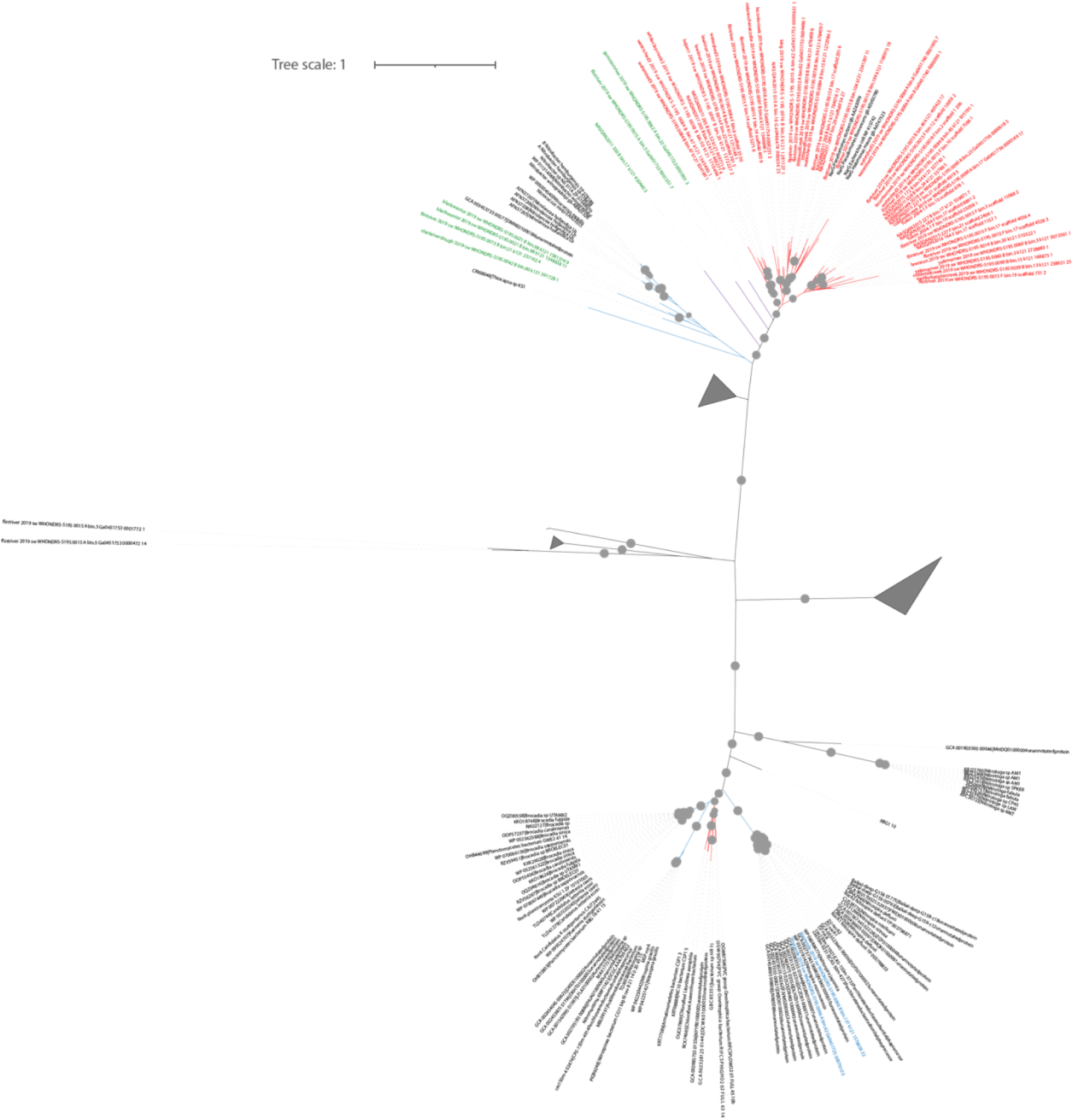
Phylogenetic analysis of nitrite oxidoreductase (nxr) and nitrate reductase (nar) is shown. GROWdb sequences colored by putative substrate, with nitrate (red), nitrite (blue), and unknown (green), while reference sequences are shown in black text. Bootstraps >80 are shown with closed grey circles at nodes. Tree files are available on Zenodo (https://doi.org/10.5281/zenodo.8173287).

**Extended Data Fig. 7:**
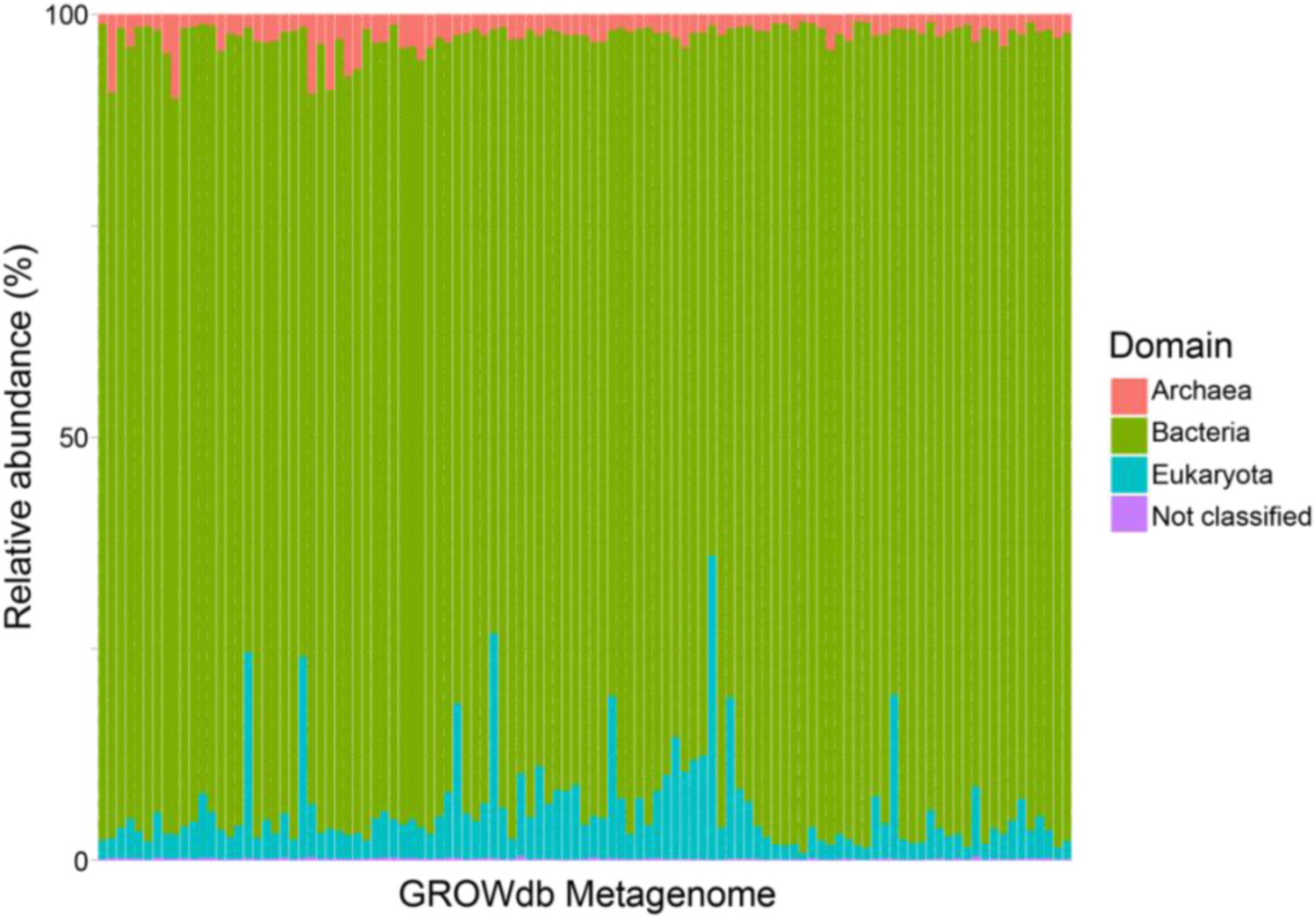
Stacked bar chart of the singleM profiles of GROWdb metagenomic reads, with bars colored by domain. By domain, the most reads are assigned to the Bacteria (mean=91.1%), followed by Eukaryota (mean=6.1%), Archaea (mean=2.6%), and Unknown (mean=0.2%).

**Extended Data Fig. 8:**
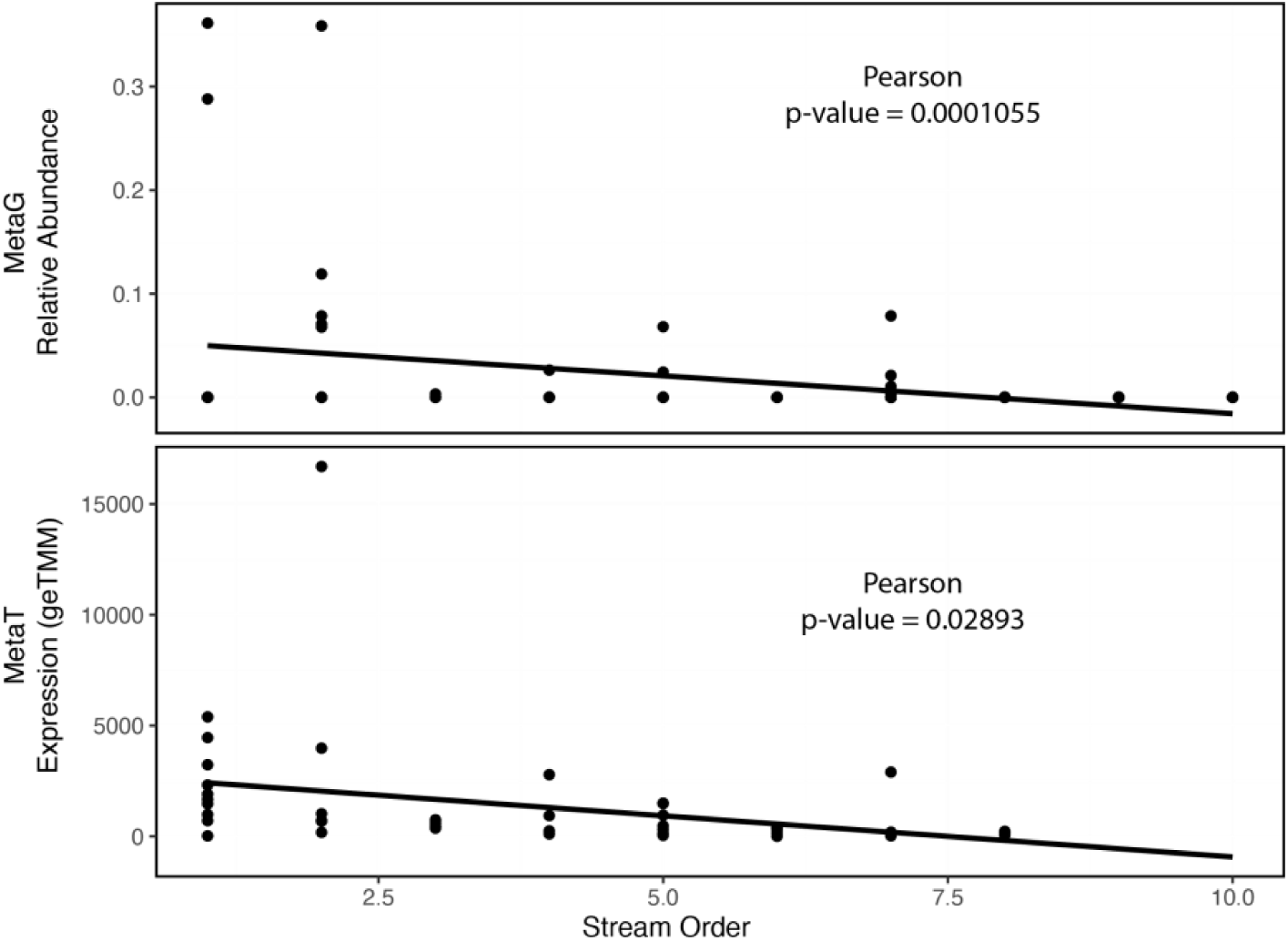
Correlations of Patescibacteria relative abundance (metagenomics, top) and expression (metatranscriptomics, bottom) with stream order.

**Extended Data Fig. 9:**
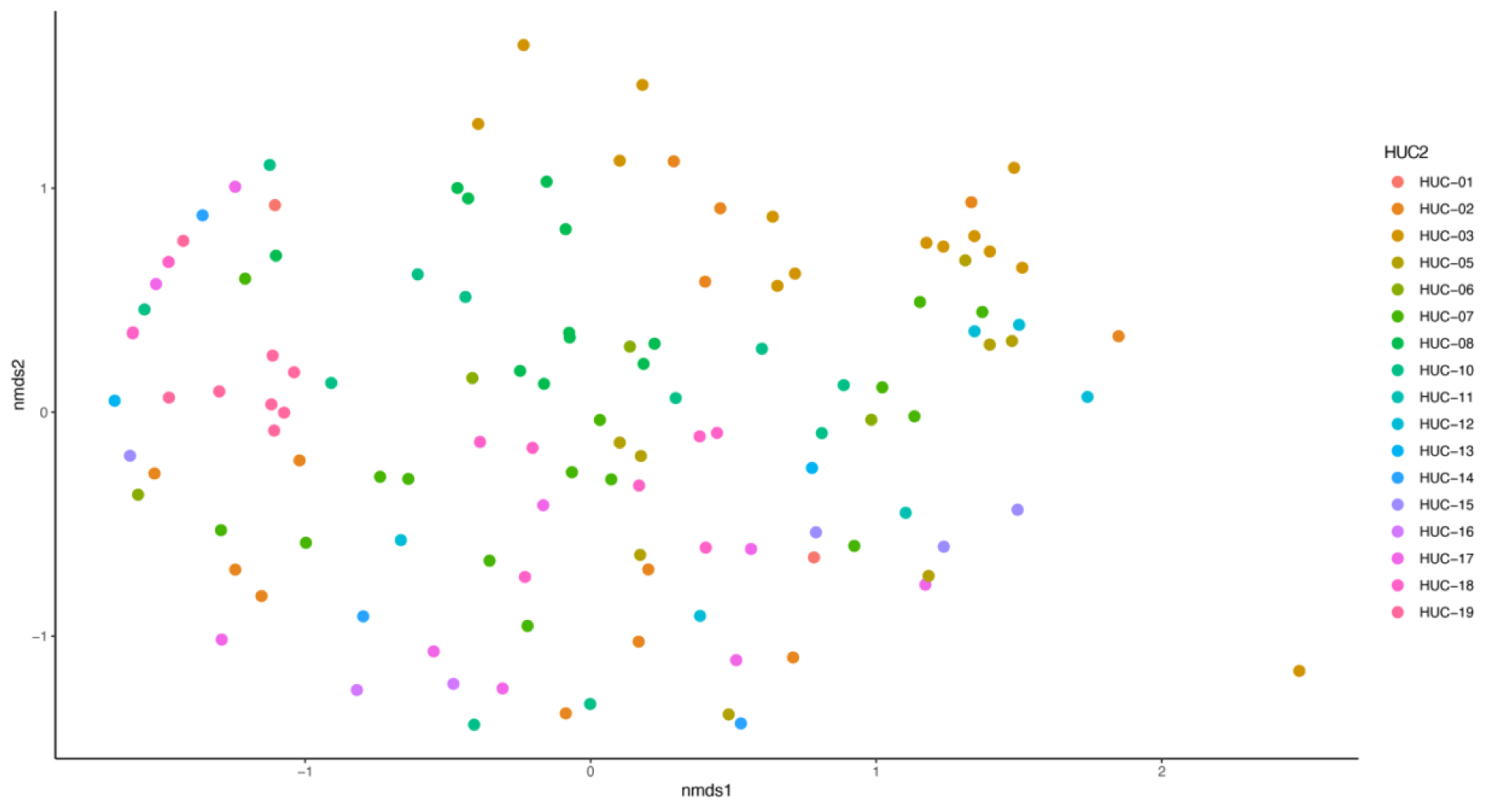
Microbial community diversity was significantly structured by hydrologic unit 2 (HUC2) region as depicted by non-metric multidimensional scaling of genome resolved metagenomic Bray-Curtis distances.

## Extended Data Files

**Extended data file 1:** Excel file.

Tab 1) Table summary of river microbiome studies detailing title, authors, and microbiome methods used. Highlighted studies are genome-resolved multi-omics like GROWdb, with only 7 rivers surveyed.

Tab 2) Metagenome and metatranscriptome information.

Tab 3) Geospatial data (n=287 variables) for each sample.

Tab 4) List of datasets included in geospatial analysis.

**Extended data file 2:** Excel file.

Tab 1) Table summary of GROWdb MAGs including taxonomy and quality information.

Tab 2) Table summary of GROWdb novelty assigned by GTDB-Tk.

Tab 3) GROWdb MAG relative abundance across metagenomes.

Tab 4) Core analysis of GROWdb genera including metagenomic and metatranscriptomic occupancy and mean abundance/expression.

**Extended data file 3:** Excel file.

Tab 1) Annotation summary of GROWdb MAGs from DRAM.

Tab 2) GROWdb whole-genome trait calls.

Tab 3) Functional genes used to make whole-genome trait calls.

Tab 4) Output of Resistance Gene Identifier containing list of predicted ARGs Tab 5) Metatranscriptomic expression of identified ARGs by category.

